# Ammonia-mediated Conformational Dynamics in SCAP for SREBP Activation

**DOI:** 10.1101/2025.05.29.656818

**Authors:** Kevin C. Chan, Zoe Li, Yan Yang, Deliang Guo, Xiaolin Cheng

**Affiliations:** Division of Medicinal Chemistry and Pharmacognosy, College of Pharmacy, The Ohio State University, Columbus, OH 43210, USA; Department of Biosciences and Bioinformatics, School of Science, Xi’an Jiaotong-Liverpool University, Suzhou, China; Department of Radiation Oncology, Ohio State Comprehensive Cancer Center, Arthur G. James Cancer Hospital and Richard J. Solove Research Institute, and College of Medicine at The Ohio State University, Columbus, OH 43210, USA; Center for Cancer Metabolism, James Comprehensive Cancer Center at The Ohio State University, Columbus, OH 43210, USA; Translational Data Analytics Institute (TDAI), The Ohio State University, Columbus, OH 43210, USA

## Abstract

Cholesterol homeostasis is regulated by the sterol regulatory element-binding protein (SREBP) pathway, with the membrane-embedded SREBP-cleavage-activating protein (SCAP) and insulin-inducible gene protein (Insig)-1/2 acting as sterol sensors. Our previous studies unveiled a critical role of ammonia in activating the SREBP pathway. Here, we performed extensive molecular dynamics (MD) simulations to elucidate the structural mechanisms through which ammonia binding triggers SCAP/Insig dissociation and subsequently activate SREBP. The SCAP S4 helix exists in two distinct conformations: a partially unfolded S4 and an intact S4; the transition between these states may constitute a crucial aspect of SCAP activation. We first studied the binding of ammonia to SCAP in different S4 conformational states. We then explored how the binding of ammonia or 25-hydroxycholesterol (25-HC) affects the conformational transition of S4 using targeted MD simulations. Finally, we performed comparative analysis of the 25-HC-bound inactive SCAP and the ammonia-bound active SCAP. Collectively, our findings highlight the importance of S4 helix conformation in activating the SCAP protein. Our simulations suggest that ammonia and 25-HC act as an agonist and antagonist of SCAP activation, respectively. Specifically, ammonia binding facilitates the transition of S4 from the unfolded to the straight conformation, triggering subsequent conformational changes in the transmembrane domain of SCAP. These changes eventually promote the SCAP/Insig dissociation and alter the position of the MELADL motif at the membrane surface.

**Significant Statement:** Cellular cholesterol homeostasis is tightly regulated by the SREBP pathway, which relies on the membrane proteins SCAP and Insig to sense sterol levels. While ammonia has recently been identified as a novel activator of this pathway, its structural mechanism of action remained unknown. Here, extensive molecular dynamics simulations reveal that ammonia binding promotes a critical conformational transition in the SCAP S4 helix, facilitating SCAP activation and its dissociation from Insig. Our findings show that ammonia and cholesterol act as opposing modulators—agonist and antagonist—of SCAP activity. This work uncovers a previously unrecognized structural switch in SCAP and provides new molecular insight into small-molecule regulation of cholesterol sensing.

## Introduction

Cholesterol plays a pivotal role in cellular physiology, serving not only as an essential component of cell membranes but also as a regulator of membrane integrity and fluidity. Cholesterol homeostasis is primarily regulated by the sterol regulatory element-binding protein (SREBP) pathway through a negative feedback mechanism.^1^ SREBP belongs to a family of transcription factors comprising three isoforms: SREBP-1a, −1c and −2. Among these, SREBP-1c regulates fatty acid synthesis, SREBP-2 is crucial for cholesterol synthesis and uptake, and SREBP-1a oversees both processes.^2^ SREBPs are synthesized in the endoplasmic reticulum (ER) as inactive precursors and require precise activation mechanisms to function.^2,3^ This activation involves a complex, multistep translocation process mediated by the SREBP-cleavage activating protein (SCAP). Structurally, SREBPs consist of an amino (N)-terminal transcription factor domain and a carboxyl (C)-terminal regulatory domain, while SCAP comprises a cytosolic C-terminal regulatory domain constitutively bound to the C-terminal domain of SREBP. The N-terminal domain of SCAP consists of eight transmembrane helices, with helices S2-S6 forming the sterol sensing domain (SSD).^4^ The dynamic interactions and structural transitions within this domain, particularly in response to cholesterol concentration fluctuations, are central to the regulation of SREBP activity, and thus cholesterol homeostasis.

SCAP fulfils dual roles as both a sterol sensor and an escort protein in the activation of SREBPs.^5^ In a sterol-rich environment, the SREBP/SCAP complex is retained in the ER membrane with the N-terminal domain of SCAP in complex with Insig-1 or Insig-2 (insulin-induced genes). This association is stabilized by cholesterol.^4^ Previous studies have shown that 25-hydroxycholesterol (25-HC), a cholesterol derivative, binds at the interface between Insig and SCAP with an even higher affinity.^6^ Upon sterol depletion, the SREBP/SCAP complex dissociates from Insig, and is then transported from the ER to the Golgi apparatus, which is facilitated by the interaction of SCAP with the coat protein complex II (COPII). Once in the Golgi, SREBP undergoes sequential cleavage by site-1 protease (S1P) and site-2 protease (S2P). This process releases the transcription factor domain, enabling its translocation into the nucleus to activate genes for lipid synthesis and uptake.

Recent studies have shown that a mere reduction in cholesterol is insufficient to activate SREBP given limited glucose availability.^7,8^ This is because glucose promotes the N-glycosylation of SCAP, enhancing its stability and reducing its susceptibility to proteasomal degradation. However, the presence of glucose alone doesn’t guarantee SREBP activation, especially under conditions where glutamine levels are low. In such scenarios, even with a low abundance of cholesterol and adequate glucose, SREBP remains inactive. This metabolic puzzle was solved with the discovery of an unexpected role of glutamine-released ammonia in SREBP activation.^9^ Ammonia is found to be essential for stimulating the dissociation of N-glycosylated SCAP from Insig, thereby initiating SREBP activation. Therefore, ammonia emerges as a critical signaling molecule, stimulating lipogenesis through the SREBP pathway. Nonetheless, the detailed structural mechanism through which ammonia promotes the dissociation of the SCAP/Insig complex remains elusive. Understanding this mechanism could provide significant insights into the regulation of lipid metabolism, potentially leading to novel therapeutic strategies for metabolic disorders and cancers.

The structure of the SCAP/Insig complex in its inactive state has been determined using cryo-electron microscopy (cryo-EM) (PDB ID: 6M49).^4^ It revealed notable structural features in which the S4 helix of SCAP is unfolded to accommodate the binding of 25-HC. In contrast, the corresponding helix in other SSD-containing proteins is typically intact.^10,11^ Yan *et al.* have hypothesized that in the absence of 25-HC, the S4 helix of SCAP might adopt an intact conformation, leading to the activation of SCAP.^4^ To elucidate the role of ammonia in SCAP activation, we employed co-solvent molecular dynamics (MD) simulations to identify a potential ammonia binding site in SCAP, located between the S3 and S6 helices. Ammonia appears to interact with this site through coordination with three key residues: D428, S326 and S330.^9^

In this study, we explore the structural transitions between the straight and bent conformations of the SCAP S4 helix, with a specific focus on their coupling with NH_4_^+^ binding. We first systematically map the ammonia distribution across various SCAP conformations. Subsequently, targeted molecular dynamics (TMD) simulations are employed to investigate the transitions of the SCAP S4 helix between its bent and straight conformations under various conditions, including the presence or absence of ammonia and different mutations. Finally, we conduct extensive unbiased MD simulations of the 25-HC-bound inactive SCAP and the ammonia-bound active SCAP to discern how local changes, such as S4 conformational transitions and ammonia binding, propagate throughout the SCAP protein. The co-solvent mapping and TMD simulations illuminate the intricate interplay between ammonia binding and the structural transition of the S4 helix and provide insights into the formation of the ammonia binding site through a conformational shift coupled with induced fit mechanism. The comparative MD simulations underscore the importance of both the S4 helix conformation and ammonia binding in orchestrating conformational changes in SCAP. These changes result in a reduction of both the interfacial contact area and polar contacts with Insig, facilitating the dissociation of the SCAP/Insig complex. Remarkably, these structural alterations also encompass a shift in the relative position of the MELADL motif in the C-terminus of the S6 helix relative to the membrane surface, suggesting potential implications for MELADL-mediated SCAP trafficking and activation. Collectively, our findings unravel the molecular mechanisms driving SCAP activation by elucidating a sequence of structural changes triggered by NH_4_^+^ binding, ultimately leading to SCAP activation.

## Methods

### Structural modeling of SCAP with a straight S4 helix

The cyro-EM structure of Insig-SCAP complex (PDB ID: 6M49) bound with 25-HC was used as the initial structure.^4^ Two unresolved loops on Insig (residue 55 to 60 and residue176 to 184) were modelled and refined using the loop refinement protocol in MODELLER V10.1.^12^ For each missing loop, a total of 10 conformations were generated, and the one with the lowest Discrete Optimized Protein Energy (DOPE) score was chosen. The S4 helix of SCAP was rebuilt using MODELLER V10.1 to replace the original unfolded region (residues 354-358) in the cryo-EM structure with a folded α-helix.^12^

### Structural modeling of SCAP with the full MELADL motif

In the cryo-EM structure, the MELADL motif was not fully resolved. To investigate the dynamics of the MELADL motif, we have added the missing residues from D451 to L456. Given that both the S4 helix conformation and the NH_4_^+^ binding are imperative for SCAP activation, we constructed two new systems: (1) Insig-SCAP complex with 25-HC bound and S4 in a bent conformation (denoted as the inactive system), and (2) Insig-SCAP in complex with NH_4_^+^ bound and S4 in a straight conformation (referred to as the active system).

### AlphaFold2 prediction

We predicted the structures of SCAP alone and the SCAP/Insig complex as well as SCAP or the MELADL motif bound to Sec proteins using LocalColabFold^13^, which combines the fast homology search of MMseqs2 with AlphaFold2^14^. The initial multiple sequence alignments (MSAs) for each protein were generated with the MMSeqs2 server^15^. We controlled the depth of the input MSAs by setting *max_msa_clusters* to 512 and *max_extra_msa* to 1024. All five neural networks were employed for non-template-based predictions. The *max_recycle* parameter was set to 3 to strike a balance between accuracy and computational efficiency, without performing the post-prediction model refinements. For each target, we selected the structure with the highest internal rank from all predicted structures for subsequent analyses. Structural alignment was performed with MatchMaker^16^ in ChimeraX^17^.

### Co-solvent mapping

To reveal ammonia binding sites in SCAP under various conditions, we utilized the co-solvent mapping (CSM) MD simulations, in which probes were added as solvent molecules. This technique dynamically explores potential binding sites or hotspots on the flexible molecular surface of SCAP.^39^ While ammonia predominantly exists in the form of ammonium ions (NH_4_^+^) under physiological conditions (pH=7), both protonated (NH_4_^+^) and deprotonated (NH_3_) forms of ammonia were simulated. CSM simulations were initiated from the cryo-EM inactive structure (bent S4)^4^ and the manually built active structure (straight S4) of SCAP-Insig. Four distinct systems were considered for NH_4_^+^ CSM simulations (Table S1): (1) *<u>CSM1</u>*: Insig-SCAP complex bound with 25-HC (bent S4); (2) *<u>CSM2</u>*: Insig-SCAP complex without 25-HC bound (bent S4); (3) *<u>CSM3</u>*: Insig-SCAP complex without 25-HC bound (straight S4); and (4) *<u>CSM4</u>*: SCAP alone (straight S4). For comparison, two systems were chosen from the above four systems for NH_3_ CSM simulations: (5) *<u>CSM5</u>*: Insig-SCAP complex bound with 25-HC (bent S4); and (6) *<u>CSM6</u>*: Insig-SCAP complex without 25-HC bound (bent S4). A complete list of all six sets of CSM simulations is summarized in Table S1.

The membrane bilayers were built using the CHARMM-GUI membrane builder, and a total of 266 hydrated palmitoyl-oleyl-phosphatidylcholine (POPC) molecules were used in each system. ^18,19^ ∼30000 TIP3P water molecules were added to each system. Na^+^ and Cl^-^ ions were added to reach an ionic concentration of 150 mM.^20^ To identify potential binding sites, either 1000 molecules of NH_3_ or 600 molecules of NH_4_^+^ were added to the system with a box size of 127Å × 127Å × 116 Å, yielding a final concentration of 1M of NH_3_ or 0.6M of NH_4_^+^. The force fields for proteins, lipids and ions were the CHARMM36 force field, while the ligand (25-HC) was parameterized with SwissParam and the co-solvent (NH_3_ or NH_4_^+^) employed the CGenFF force field.^21–23^ The system was initially prepared using Gromacs 2020 and converted to AMBER formats using ParmEd.^24–26^ All co-solvent mapping simulations were conducted at 310K with the Langevin thermostat using AMBER18.^24,26^ A semi-isotropic Monte Carlo barostat was employed to maintain the pressure at 1 bar. The integration time step was 2 fs. Non-bonded interactions were calculated with the particle mesh Ewald (PME) method with a 12 Å cutoff ^27^. The SHAKE algorithm was used for all bonds containing hydrogen ^28^.

All simulations underwent an initial energy minimization using the steepest-descent method, with a maximum of 50,000 steps. Following energy minimization, a 500 ps equilibration simulation was conducted with position restraints applied to the protein, lipids, and ligands. Subsequently, six 1 ns simulations were performed consecutively, gradually reducing the strength of the positional restraints. The unrestrained production simulations were started from the final frame of these restrained simulations. Finally, six sets of simulations (Table S1), each with five independent replicas, were conducted. This resulted in a total of 30 simulations, each with a duration of 100 ns.

### Molecular dynamics simulations

MD simulations of the Insig-SCAP complex embedded in a POPC bilayer were prepared as described above. Five simulation systems were constructed (Table S2): (1) *<u>MD1</u>*: Insig-SCAP complex bound with NH_4_^+^ (bent S4 helix), (2) *<u>MD2</u>*: Insig-SCAP complex bound with NH_4_^+^ (straight S4), (3) *<u>MD3</u>*: Insig-SCAP complex bound with 25-HC (bent S4), (4) *<u>MD4</u>*: Insig-SCAP complex without 25-HC or NH_4_^+^ bound (straight S4), and (5) *<u>MD5</u>*: Insig-SCAP complex without 25-HC or NH_4_^+^ bound (bent S4). MD1 and MD2 were initiated from the final frames of the corresponding co-solvent mapping (CSM) simulations. MD3 was initialized from the cryo-EM structure of the Insig-SCAP complex (PDB ID: 6M49). MD4 was generated by removing 25-HC from the same structure, while MD5 was prepared by additionally replacing the S4 helix with a modeled straight conformation, as described in the Structural Modeling section.

Each system was solvated with ∼34,000 TIP3P water molecules and neutralized with 150 mM NaCl. The same force fields were used for proteins, lipids, ions, and ligands as described above. All simulations were prepared and executed using GROMACS 2020.^25^ Simulations were conducted in the NPT ensemble at 310 K using the v-rescaling thermostat,^29^ with protein, membrane and ligand coupled separately from water and ions with a relaxation time constant of 0.1 ps. Pressure was maintained at 1 bar using the Parrinello-Rahman barostat with a coupling constant of 0.2 ps and an isothermal compressibility of 4.5 × 10^-5^ bar ^-1^.^30^ Semi-isotropic pressure coupling was applied with the x and y directions coupled independently from the z direction. Long-range electrostatic interactions were calculated using the particle mesh Ewald method with a real-space cutoff of 14 Å.^27^ Van der Waals interactions were truncated at 14 Å, and the neighbor list was updated every 20 steps using the Verlet cutoff scheme. All bonds were constrained with the LINCS algorithm.^31^ The integration time step was set at 2 fs. Energy minimization and equilibration protocols were the same as those in the CSM simulations as described above. Multiple μs-long production runs, starting from the last frame of the corresponding equilibration simulation, were performed for each system.

### Targeted molecular dynamics (TMD) simulations

The wild type Insig-SCAP complex embedded in a POPC bilayer was prepared as described above. Two SCAP point mutants, Y298C and D428A, were generated using UCSF Chimera^32^ and similarly embedded in the bilayer using identical protocols. All TMD simulations were performed using AMBER18.^24,26^ TMD simulations were conducted in both forward and backward directions to investigate conformational transitions of the SCAP S4 helix. In forward simulations, the initial structure corresponded to the inactive state of SCAP, in which the S4 helix adopts a bent (unfolded) conformation based on cryo-EM data, while the target structure was the active conformation with a straight (folded) S4 helix. Conversely, in backward simulations, the straight S4 conformation served as the starting structure, and the bent conformation was used as the target.

All minimization and equilibration steps were identical to those described for the previous systems. During the production phase of TMD simulations, the steered MD option was enabled with the path mode set to *spline*. Steering forces were applied exclusively to the backbone atoms of the S4 helix to drive the conformational transition. For each system, a range of harmonic force constants from 20 to 150 kcal×mol^-1^×Å^-2^ was tested (see Figure S2 for details). For each force constant, 10 to 60 independent simulations were conducted, each with a duration of 10 ns, resulting in over 1,500 TMD simulations and a cumulative simulation time exceeding 15 μs.

To assess the kinetics and success rate of the conformational transition, we monitored the presence of 11 potential α-helical hydrogen bonds spanning residues 351 to 361, between backbone oxygen and nitrogen atoms of residue pairs i and i+4 (e.g. 351-355 to 361-365) (Figure S3). A hydrogen bond was considered present if the donor-acceptor distance was within 4 Å. A successful bent-to-straight transition was defined as the formation of at least 10 out of 11 hydrogen bonds, sustained for a minimum duration of 500 ps. The same criteria, applied in reverse, defined a successful straight-to-bent transition. Activation time was recorded as the time at which the transition criteria were first satisfied. The success rate was calculated as the fraction of simulations that met these criteria within the 10 ns simulation window.

As expected, lower force constants generally resulted in longer activation times and lower success rates, while higher force constants consistently induced transitions within the 10 ns timeframe. To quantify the difficulty of the S4 conformational transition, we introduced the metric of effective stiffness, defined as the harmonic force constant required to achieve 50% normalized activation efficiency. Activation times were first normalized by dividing each value by the maximum activation time observed across all force constants. A sigmoid function was then fitted to the normalized activation time (y) as a function of the applied force constant (x).

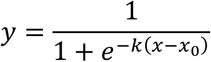

where *k* represents the slope and *x*_0_is the midpoint of the curve, corresponding to the force constant at which the normalized activation time reaches 50% of its maximal value. The *x*_0_ value is taken as the effective stiffness of the system.

### Data analysis

Structural visualization was performed using VMD and Chimera.^32,33^ Distances between S326, S330, and D428, as well as four residue pairs within the unfolded region of the S4 helix, were computed using VMD.^33^ Specifically, the distances between S326, S330 and D428 were measured between the OG atoms of the serine residues and the COM of the OD1 and OD2 atoms of D428. Distances between residue pairs within the unfolded region of S4 helix were measured using the backbone oxygen and nitrogen atoms.

The tilt angle of each helix was determined by calculating the angle between the principal axis of the helix and the z-axis. The kink angle of the S5 or S6 helix was defined as the angle between the principal axes of the upper and lower halves of each helix. To quantify interfacial interactions, the contact surface area was computed by measuring the solvent accessible surface area of TM3 and TM4 of Insig and S2, S4, and S5 helices of SCAP using the *sasa* function in VMD. The number of polar contacts was computed using the *measure contact* function in VMD with a 4 Å cutoff. Only polar or charged residues facing the interface were considered in this analysis. The position of the MELADL motif was tracked by measuring the z coordinates of the Cα atoms of the six motif residues relative to the z-coordinates of the lipid phosphate atoms. All plots were generated using Matplotlib.^34^ In box plots, the box extends from the first to the third quartile (interquartile range, IQR), with a horizontal line indicating the median. Whiskers extend from the box to the most extreme data points within 1.5× IQR from the box, and data points beyond this range are shown as fliers (outliers).

## Result and Discussion

### Structural modeling of SCAP with a fully folded S4 helix

A distinctive feature of the recent cryo-EM structure of the SCAP/Insig complex is the partially unwound S4 helix, forming a kink in its middle from residues Val355 to Leu358. This kink shifts the bottom half of the S4 helix away from the SCAP/Insig interaction interface, allowing accommodation for 25-HC. This unwound S4 conformation differs from that in other SSD-containing proteins, such as NPC1 and Ptch1, where the S4 helix typically remains intact.^10,11^ Such structural transitions in the S4 helix have been hypothesized to be pivotal in SCAP activation, as evidenced by the effects of well-characterized SCAP mutations, including D428A and Y298C in pull-down and SREBP activation assays.^35–37^ To explore SCAP’s active state without 25-HC or Insig, we manually modeled its structure with a straightened S4 helix (Figure 1a). Additionally, for comparison purpose, we predicted the structures of SCAP alone, presumably in an active state, and the SCAP-Insig complex in an inactive state using AlphaFold2 (AF2)^14^ and AF2 multimer^38^ (Figure 1b). Our manually modeled SCAP structure aligns well with that of the ‘active’ SCAP predicted by AF2, with a root-mean-square-deviation (RMSD) of 1 Å for the SSD domain, supporting the notion of the S4 being fully folded in the active state.

**Figure 1.**
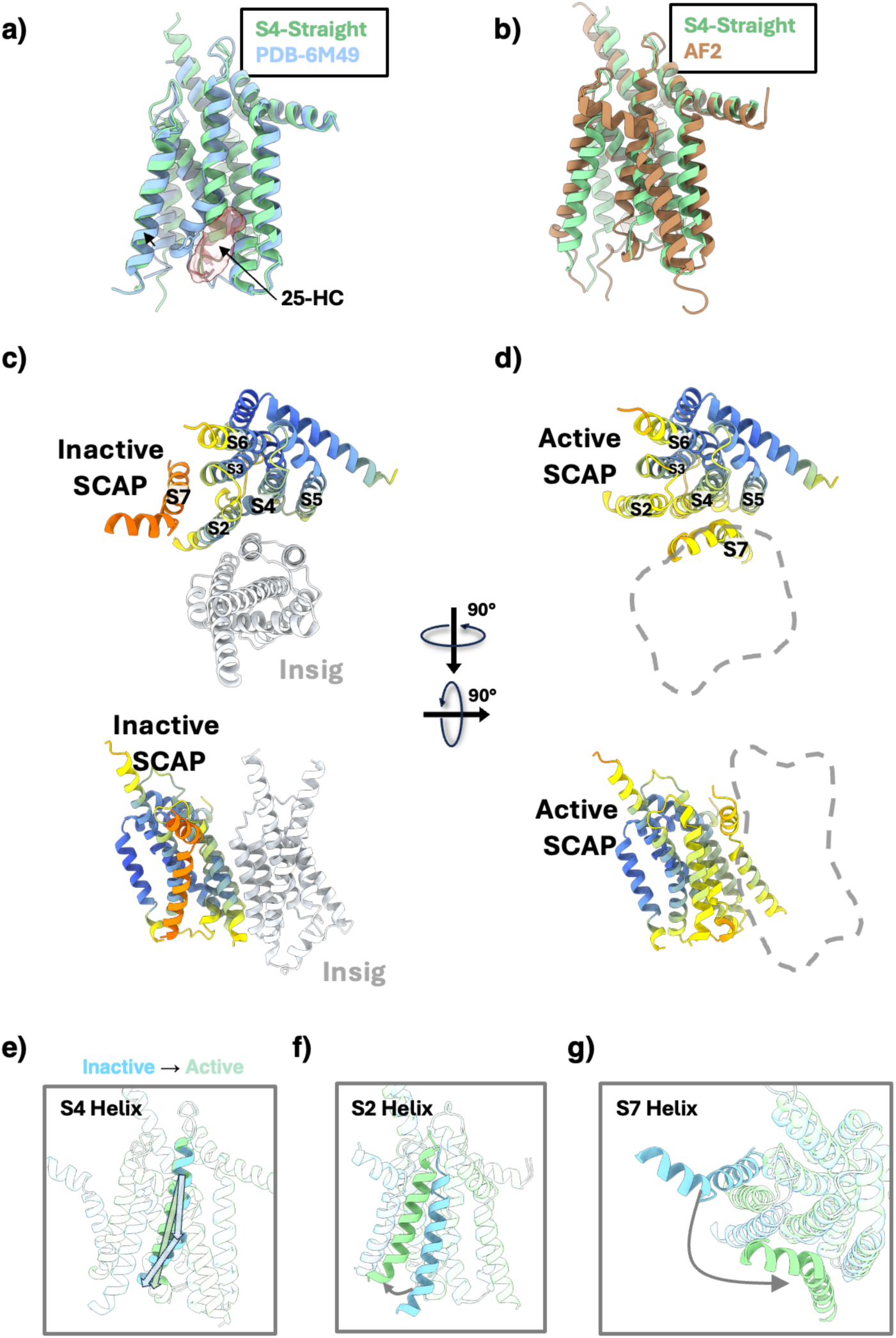
Superimposition of SCAP structures with variations in S4 helix conformation. SCAP structure with a modeled straight S4 helix is aligned to (a) structure with a bent S4 helix (PDB ID: 6M49) or with (b) structure with a straight S4 helix predicted by AF2. AF2-predicted structures of (c) inactive and (d) active SCAP are colored based on pLDDT scores. Blue indicates the highest confidence, while orange indicates lower confidence. (e-g) Three major structural differences between the inactive (blue) and active (green) SCAP are shown.

A comparison of the AF-predicted structures of inactive and active SCAP unveiled three noteworthy differences in SCAP. Firstly, in line with our hypothesis, a bent S4 was observed in the inactive state as opposed to a straight S4 in the active state (Figure 1c), supporting the idea that the conformation of the S4 helix is a crucial determinant of SCAP’s functional state. It is important to note that the AF2 model was not informed about the presence of 25-HC at the protein interface. Therefore, the predicted bent S4 in the SCAP/Insig complex is solely attributed to the presence of the Insig protein. Secondly, the transition of SCAP from an inactive to an active state involves the displacement of the S2 helix from the 25-HC binding site, resulting in the disruption of a hydrogen bond between Tyr298 and Leu358, previously identified as stabilizing the bent S4 conformation ^4^ (Figure 1d). This observation aligns with the GOF Y298C mutation.

Lastly, in the active SCAP, the S7 helix was predicted with high confidence to be positioned next to S4 and S5, thereby sterically blocking Insig binding (Figure 1b). In contrast, in the inactive SCAP, the S7 helix appears to move away from the Insig binding interface with very low prediction confidence (Figure 1b,e), suggesting a dynamic role of the S7 helix in SCAP activation.

### NH4^+^ binding in SCAP

The CSM simulations reveal that NH_4_^+^ clustered near the S6 helix in all three CSM systems lacking 25-HC (CSM2, CSM3 and CSM4), suggesting a potential binding site despite the distinct conformation of the S4 helix (Figure 2d-f). This site was absent in CSM1 with 25-HC bound (Figure 2c), where most NH_4_^+^ ions were concentrated on the top and bottom plane of the protein complex, indicating competitive binding between NH_4_^+^ and 25-HC. The absence of 25-HC creates space at the SCAP/Insig interface, allowing NH_4_^+^ to reach the binding site. NH_3_ molecules, however, were found outside the protein complex independent of the presence of 25-HC (Figure S1).

**Figure 2.**
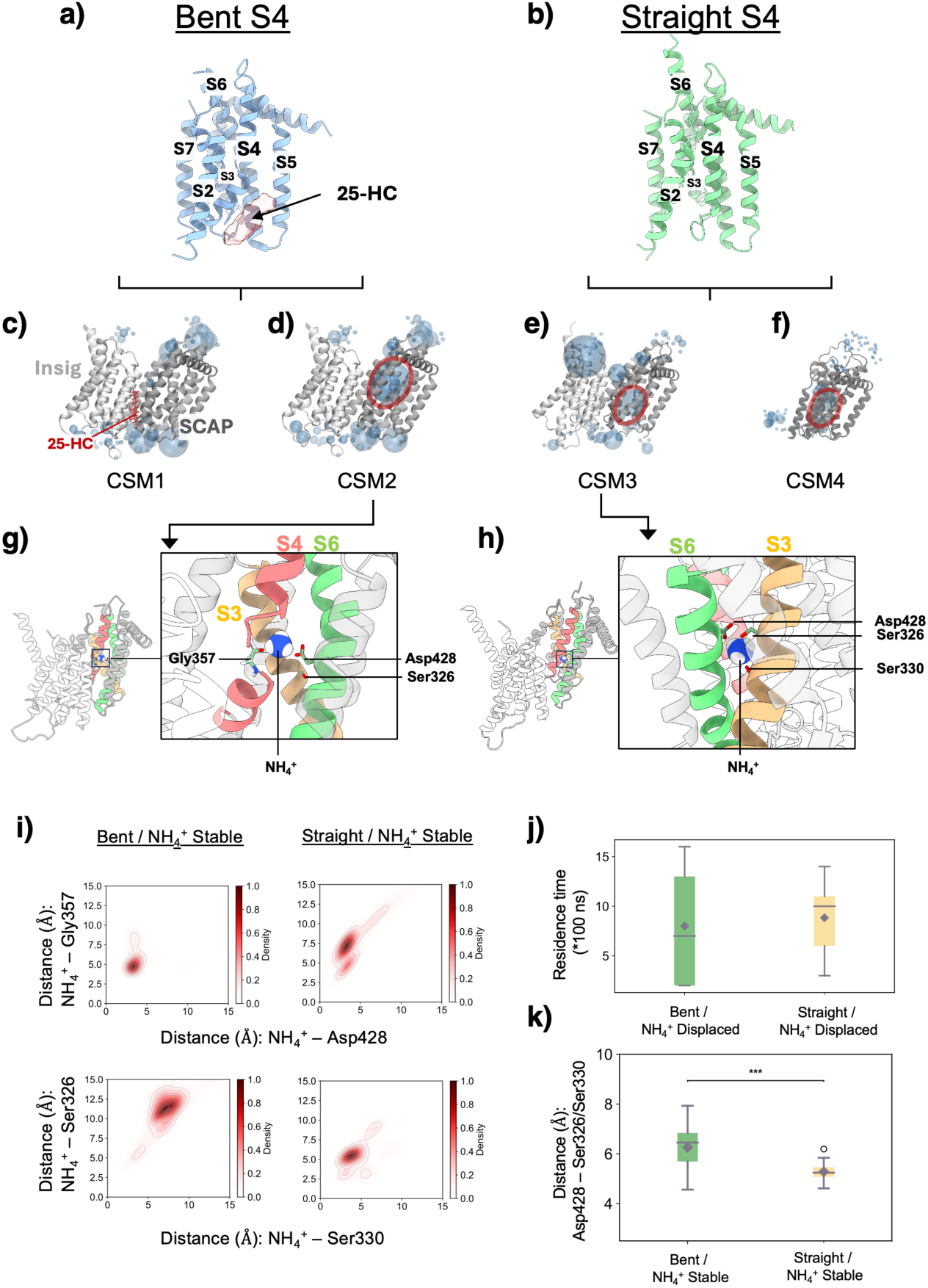
NH_4_^+^ distributions in the SCAP protein. Structures with S4 helix in (a) bent and (b) straight conformations are shown. For CSM simulations (c-f), blue spheres represent the distributions of co-solvents (NH_4_^+^ molecules) near the protein complex, with their radii proportional to the density averaged over the simulation trajectories. Red circle highlights the specific site where NH_4_^+^ binds to the SCAP protein. CSM simulation snapshots of NH_4_^+^ binding site in SCAP (g) with a bent S4 and (h) with a straight S4. NH_4_^+^ is shown as a blue sphere. Key residues are represented as licorices. Hydrogen bond occupancies are computed by averaging across multiple trajectories. (i) Density plots of distance between the NH_4_^+^ molecule and Asp428, Gly357, Ser326, and Ser330 measured from MD simulations. (j) Residence time of NH_4_^+^ molecule measured from MD simulations in which NH_4_^+^ eventually disassociates from the site. (j) Distances between Asp428 and Ser326/Ser330 in SCAP measured from MD simulations in which the NH_4_^+^ molecule stably bind to the SCAP.

**Figure 3.**
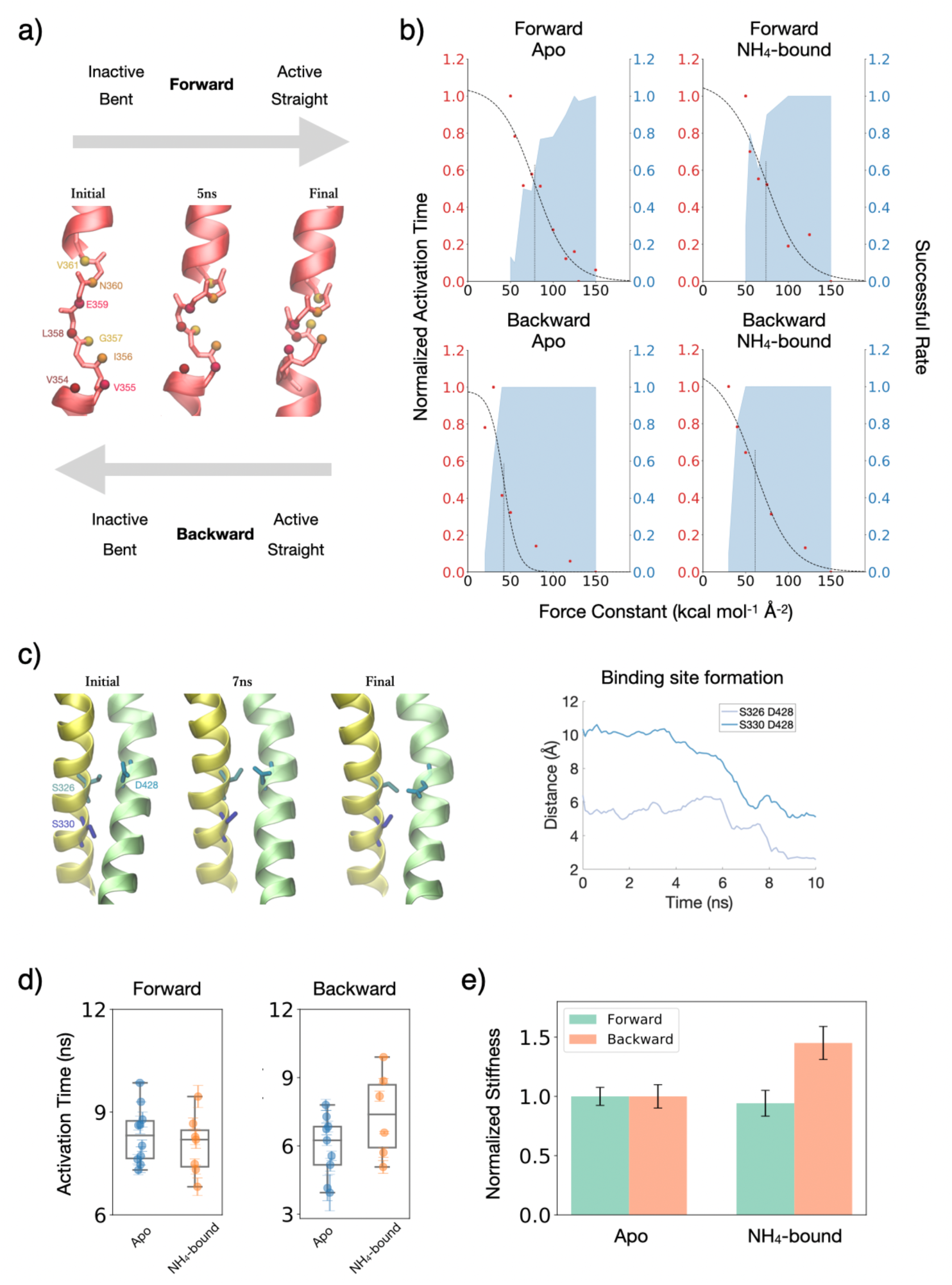
TMD simulations of structural transitions between the two S4 helix conformations. (a) Schematics illustrating the forward transition (S4 transitions from a bent to a straight conformation) and the backward transition (S4 transitions from a straight to a bent conformation). (b) Activation times and success rates as a function of force constant in the TMD simulations for both apo and NH_4_^+^-bound SCAP. (c) Formation of a binding site between helices S3 and S6 during the TMD simulations. (d) Comparison of activation times between apo and NH_4_^+^-bound SCAP for both forward and backward transitions. (e) Comparison of structural stiffness between apo and NH_4_^+^-bound SCAP for both forward and backward transitions.

**Figure 4.**
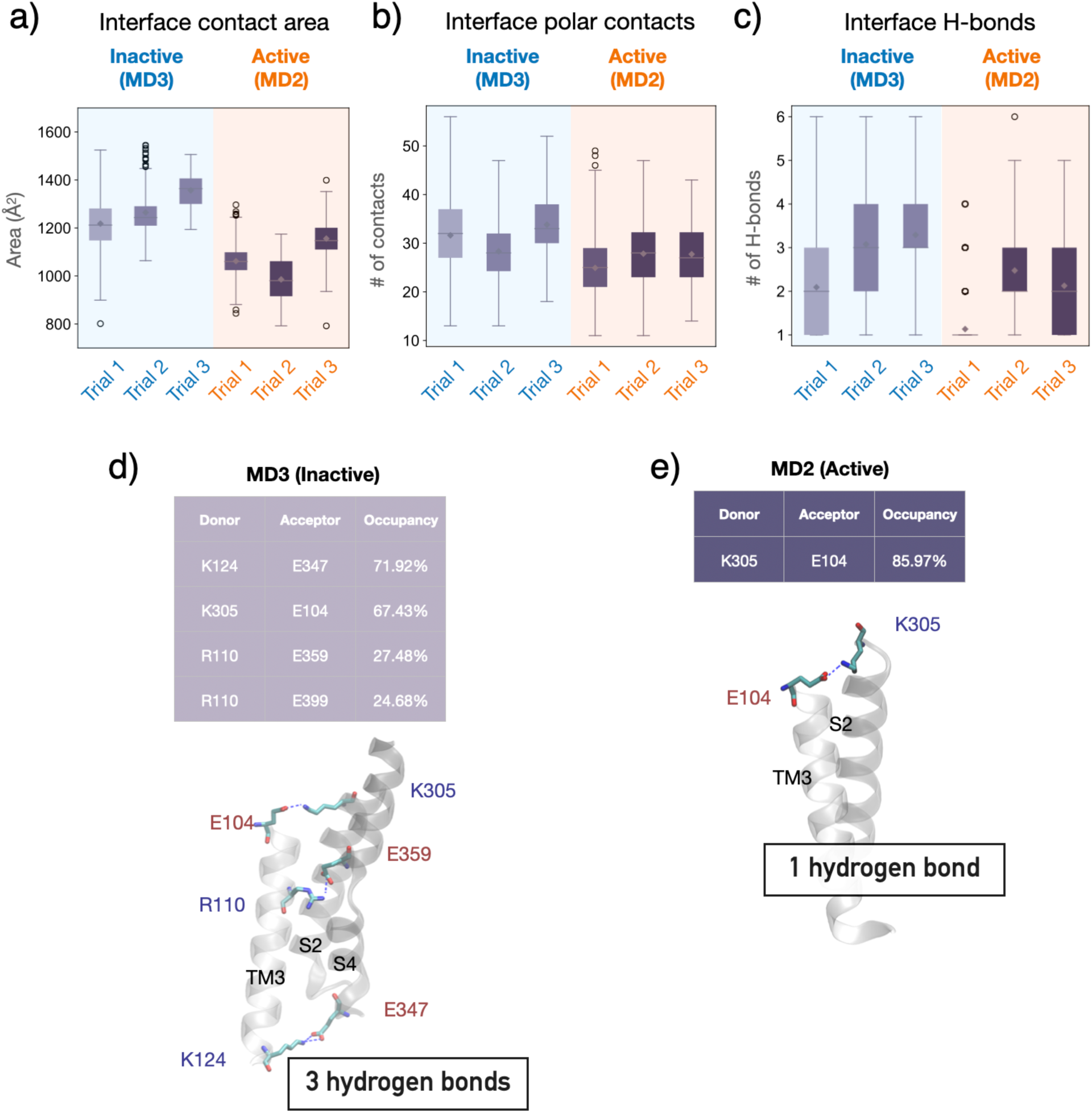
Interactions between SCAP and Insig during the simulations of inactive (MD3) and active (MD2) systems. (a-c) Comparison of interfacial contact areas, polar contacts, and hydrogen bonds between inactive and active systems. (d) Snapshot from MD3, illustrating three hydrogen bonds at the interface in the inactive SCAP/Insig complex. (e) Snapshot from MD2, showing a single hydrogen bond at the interface of the active SCAP/Insig complex.

The key difference between CSM2 and CSM3 lies in the conformation of the SCAP S4 helix: bent in System 2 vs. straight in System 3. Inspection of the NH_4_^+^ distributions in the simulation trajectories revealed notable differences in NH_4_^+^ binding sites between these systems. In CSM2, the unfolded region of the S4 helix exposes the backbone carbonyl oxygen atom of Gly357, attracting positively charged NH_4_^+^ (Figure 2g). NH_4_^+^ binding to this site was consistently observed in all five trajectories, resulting in an overall occupancy of 40.4%. Conversely, the straight conformation of the S4 helix in CSM3 prevents the interaction of Gly357 with NH_4_^+^. This structural variance redirects NH_4_^+^ to a new binding site situated between the S3 and S6 helices, otherwise shielded by the bent S4 helix (Figure 2h). NH_4_^+^ binds to the site with an even higher occupancy of 51.6%, primarily coordinated by three residues: Asp428 from the S6 helix, and Ser326 and Ser330 from the S3 helix. Among them, the negatively charged side chain of Asp428 contributes the most to the binding. In both systems, a competitive binding relationship was observed between NH_4_^+^ and 25-HC. In the presence of 25-HC, the binding site appears inaccessible to NH_4_^+^ from the bulk solvent. However, in its absence, as in CSM2, NH_4_^+^ could pass through the SCAP/Insig interface, with the exposed backbone carbonyl oxygen atom of Gly357 readily accessible for NH_4_^+^ binding.

To substantiate the identified NH_4_^+^ binding sites, we performed additional MD simulations on two NH_4_^+^-bound SCAP/Insig complex structures derived from CSM2 and CSM3. These systems are denoted as MD1 (bent S4 with NH_4_^+^) and MD2 (straight S4 with NH_4_^+^), respectively. For comparison, MD simulations were also initiated on systems without NH_4_^+^, referred to as MD3 (bent S4 with 25-HC and without NH_4_^+^) and MD4 (straight S4 with neither 25-HC nor NH_4_^+^). In MD1 (bent S4), NH_4_^+^ remained in the binding site in 6 out of 10 simulations. Similarly, in MD2 (straight S4), NH_4_^+^ occupied the binding site in 5 out of 10 simulations (Figure 2i). In both cases, the observed binding events suggest a moderate to sustained binding affinity, as inferred from the residence times in simulations where NH₄⁺ eventually dissociates from the site. Notably, MD1 exhibited a broader range of NH₄⁺ residence times (Figure 2j), spanning from approximately 200 ns to 1600 ns, indicating greater variability in binding stability compared to MD2. During the MD2 simulation, NH_4_^+^ formed stable hydrogen bonds primarily with the side chains of Asp428, Ser326, and Ser330, with Asp428 contributing the most to NH_4_^+^ stabilization, as evidenced by hydrogen bond occupancy. This predicted binding site was validated by the co-immunoprecipitation assays, where the SCAP D428A mutant completely abolished NH_4_^+^ binding and SREBP-1 activation, and the S326A/S330A double mutant significantly reduced SREBP-1 activation.^9^ Intriguingly, an induced fit effect is evident for the NH_4_^+^ binding site in the S4 straight system. That is, NH_4_^+^ binding facilitated the binding site formation. Comparison of MD2 (NH_4_^+^ bound) with MD4 (no NH_4_^+^ bound) reveals a noticeable decrease in the average distances between Ser326 or Ser330 and Asp428 (Figure 2k and Table 1). However, in systems with a bent S4 helix (MD1 and MD3), NH_4_^+^ binding showed no effect on the distances between Ser326 or Ser330 and D428, regardless of the presence of 25-HC or Insig. Taken together, our MD simulations confirmed that NH_4_^+^ preferentially binds to the ‘active’ SCAP (straight S4) compared to the ‘inactive’ SCAP (bent S4). This suggests that NH_4_^+^ could act as an agonist in SCAP activation, preferentially stabilizing the active state over the inactive state of SCAP.

**Table 1.** Comparison of average distances between Ser326 or Ser330 and Asp428 in SCAP during the simulations.

| MD System | S326—D428 Distance (Å) | S330—D428 Distance (Å) |
| --- | --- | --- |
| 1 | $5.5 \pm 0.8$ | $9.8 \pm 0.7$ |
| 2 | $3.2 \pm 0.9$ | $5.2 \pm 0.6$ |
| 3 | $5.9 \pm 0.6$ | $9.8 \pm 0.7$ |
| 4 | $3.7 \pm 1.0$ | $7.1 \pm 0.7$ |
| 5 | $4.6 \pm 1.1$ | $8.5 \pm 1.5$ |

### Explore the structural transition of the S4 helix between two conformational states using targeted MD (TMD) simulations

We next performed extensive TMD simulations to explore how the binding of 25-HC or NH_4_^+^ may influence the structural transition between the bent and straight conformations of the S4 helix. We denote the transition from the bent to straight conformation as the *forward* transition, while the reverse is referred to as the *backward* transition (Figure 5a). Constructing a structure of the 25-HC-bound SCAP/Insig complex with S4 in a straight conformation proved challenging. In the presence of 25-HC, the S4 helix breaks into two segments. The tilting of the bottom half of S4 creates space for 25-HC occupancy, while a fully folded straight S4 helix would clash with the bound 25-HC. This indicates that only the inactive, bent conformation of S4 can accommodate 25-HC, suggesting that in contrast to the agonistic role of NH_4_^+^, 25-HC functions as an antagonist, favoring the inactive state of SCAP or driving the transition from an active to an inactive state.

TMD simulations were first performed on the SCAP/Insig complex structures without the binding of either NH_4_^+^ or 25-HC (apo state). These simulations employed various force constants, each with multiple independent replicates, as shown in Figure S2. To monitor the progression of the transition during TMD simulations, we measured distances between 11 pairs of backbone oxygen and nitrogen atoms within S4 helix. The criteria for defining the active state and the corresponding activation time, based on dynamic hydrogen bond networks, are detailed in the Methods section and Figure S3. In *forward* simulations, a complete transition required a minimum force constant of 50 kcal/mol/Å^2^ (Figure 5b). This transition occurred gradually, characterized by the sequential formation of backbone hydrogen bonds. Not surprisingly, the f*orward* transition coincided with the formation of the NH_4_^+^ binding site, evidenced by the convergence of Ser326 and S330 to Asp428 (Figure 5c). Conversely, the *backward* transition necessitated a lower force constant (Figure 5b), proceeding in a concerted manner with simultaneous formation of multiple backbone hydrogen bonds. Consistently, in the apo state, the *backward* transition required shorter activation times compared to the *forward* transition (Figure 5d). These results suggest that the *backward* transition occurs more readily, indicating that the straight conformation of the S4 helix is less stable than its bent counterpart in the absence of both NH_4_^+^ and 25-HC. This finding aligns with experimental data, suggesting that the SCAP activation is unlikely in the absence of NH_4_^+ 9^.

To explore the impact of NH_4_^+^ binding on the conformational transition of SCAP, we conducted TMD simulations on the NH_4_^+^-bound SCAP/Insig complex. To quantitatively compare the transition in NH_4_^+^-bound and apo systems, we introduced the metric of effective stiffness, defined as the force constant required to reduce the activation time to half of its maximum value. In the NH_4_^+^-bound system, the effective stiffness for the *forward* transition was 73.89 ± 8.54 kcal/mol/Å^2^. In contrast, the apo state exhibited a slightly increased stiffness of 78.45 ± 5.97 kcal/mol/Å^2^ for the same transition (Figure 5e). For the *backward* transition, the NH_4_^+^-bound system displayed a stiffness of 61.12 ± 5.86 kcal/mol/Å^2^, while the apo system showed a much-reduced stiffness of 42.13 ± 4.15 kcal/mol/Å^2^. Notably, at sub-saturation force constants, the NH_4_^+^-bound system demonstrated a higher probability of completing the *forward* transition compared to the apo system (Figure 5b). Overall, these TMD simulation results suggest that NH_4_^+^ binding facilitates the transition from an inactive to an active state of the S4 helix. Furthermore, the balance appears to be tipped through a preferential stabilization of the straight (active) conformation over its bent (inactive) counterpart upon NH_4_^+^ binding.

To substantiate our findings regarding the conformational transition of the S4 helix, we performed TMD simulations on two well-characterized SCAP mutants: the gain-of-function (GOF) mutant Y298C and the loss-of-function (LOF) mutant D428A (Figure S4). These mutations, along with L315F and D443N, have been extensively studied for their impact on SCAP function.^4,35,36,40^ The Y298C mutation, involving the substitution of tyrosine at position 298 with cysteine, leads to the disengagement between SCAP and Insig, indicative of a shift in balance towards the straight S4 (active) conformation. In line with this observation, our *forward* TMD simulations of the Y298C mutant revealed lower stiffness compared to the wild type in its apo form (Figure S5). Conversely, the D428A mutation, where aspartic acid at position 428 is replaced by alanine, was found to enhance the stability of the SCAP/Insig complex, suggesting a preference for the bent S4 (inactive) conformation. As expected, the *forward* transition in the D428A mutant exhibited increased stiffness compared to the wild type, while the *backward* transition showed a slight decrease in stiffness (Figure S5). Overall, our TMD simulation results are in broad agreement with experimental findings of the GOF and LOF effects in SCAP mutants. A clear correlation emerges between the conformation of the S4 helix and the functional state of SCAP, underscoring the significance of these conformational transitions in SCAP activation.

### Ammonium binding coupled with S4 conformational transition leads to structural changes in other regions of SCAP

To elucidate the structural mechanisms underlying NH_4_^+^ binding in the activation process, we extended the simulations for all MD systems, and compared the dynamic behavior between the inactive (MD3 - bent S4 with 25-HC and without NH_4_^+^ bound) and the active (MD2 - straight S4 with NH_4_^+^ and without 25-HC bound) states to shed light on its role in facilitating SCAP’s dissociation from Insig. We first quantified the interfacial contact area between SCAP and Insig in these simulations. A pronounced decrease in interfacial contact area was observed for the active system compared to the inactive system (Figure 6a). This reduction was accompanied by a minor decrease in the number of interfacial polar contacts (Figure 6b). Remarkably, the interfacial area was found to be primarily influenced by the conformational state of the S4 helix, whereas the presence of 25-HC or NH_4_^+^ did not significantly alter the interfacial area, as evidenced by the comparable areas in systems with the respective bent and straight S4 conformations (Figure S6).

**Figure 5.**
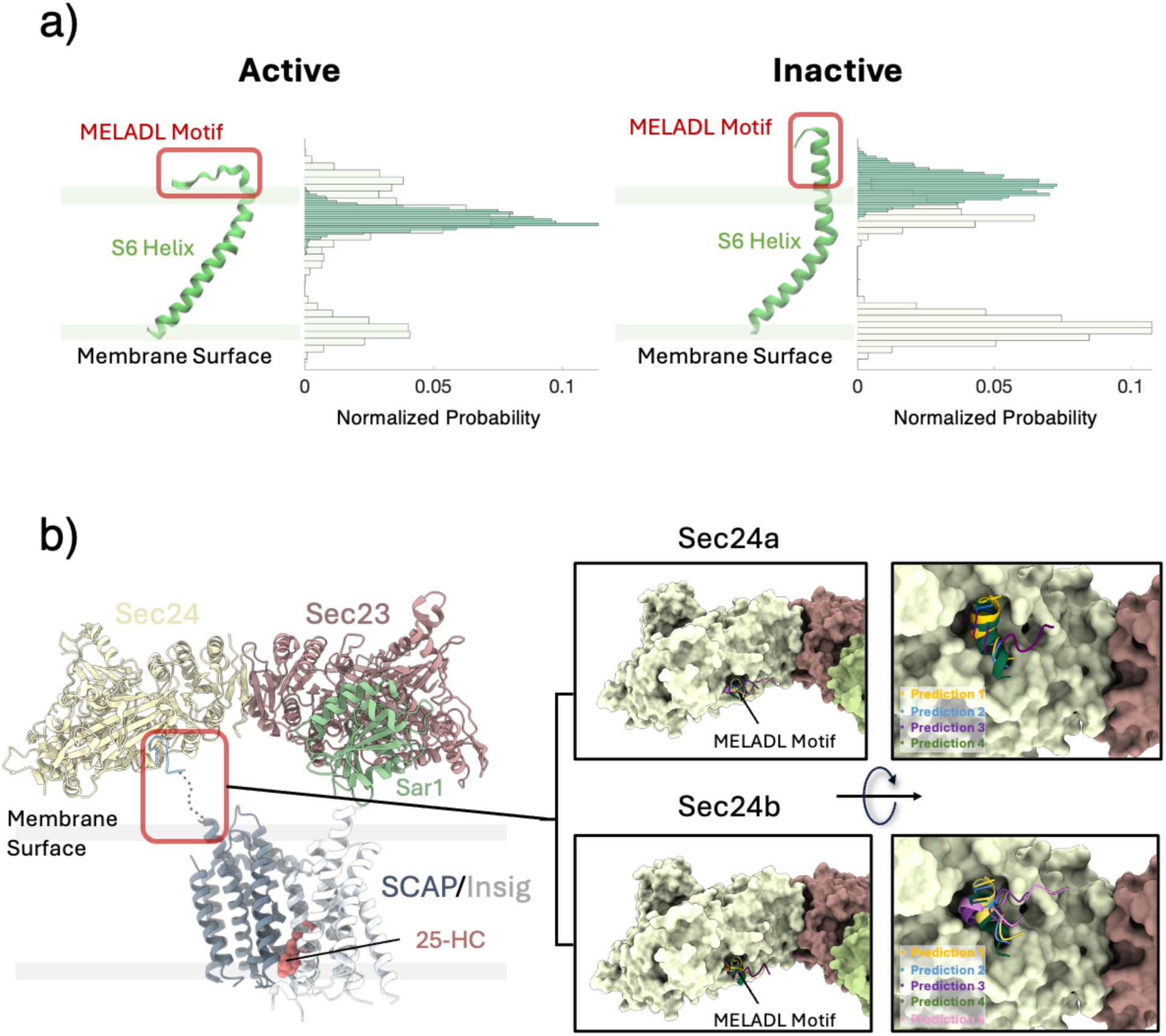
Interaction of the MELADL motif with Sec proteins. (a) Structure and position of the MELADL motif during the simulations of active (MD2) and inactive (MD3) SCAP. (b) Schematic representation of the interaction between the SCAP/Insig complex and the Sec protein complex. Enlarged views illustrate the AF2 predicted structures of MELADL motif binding to Sec24a and Sec24b proteins, highlighting the potential interaction modes.

**Figure 6.**
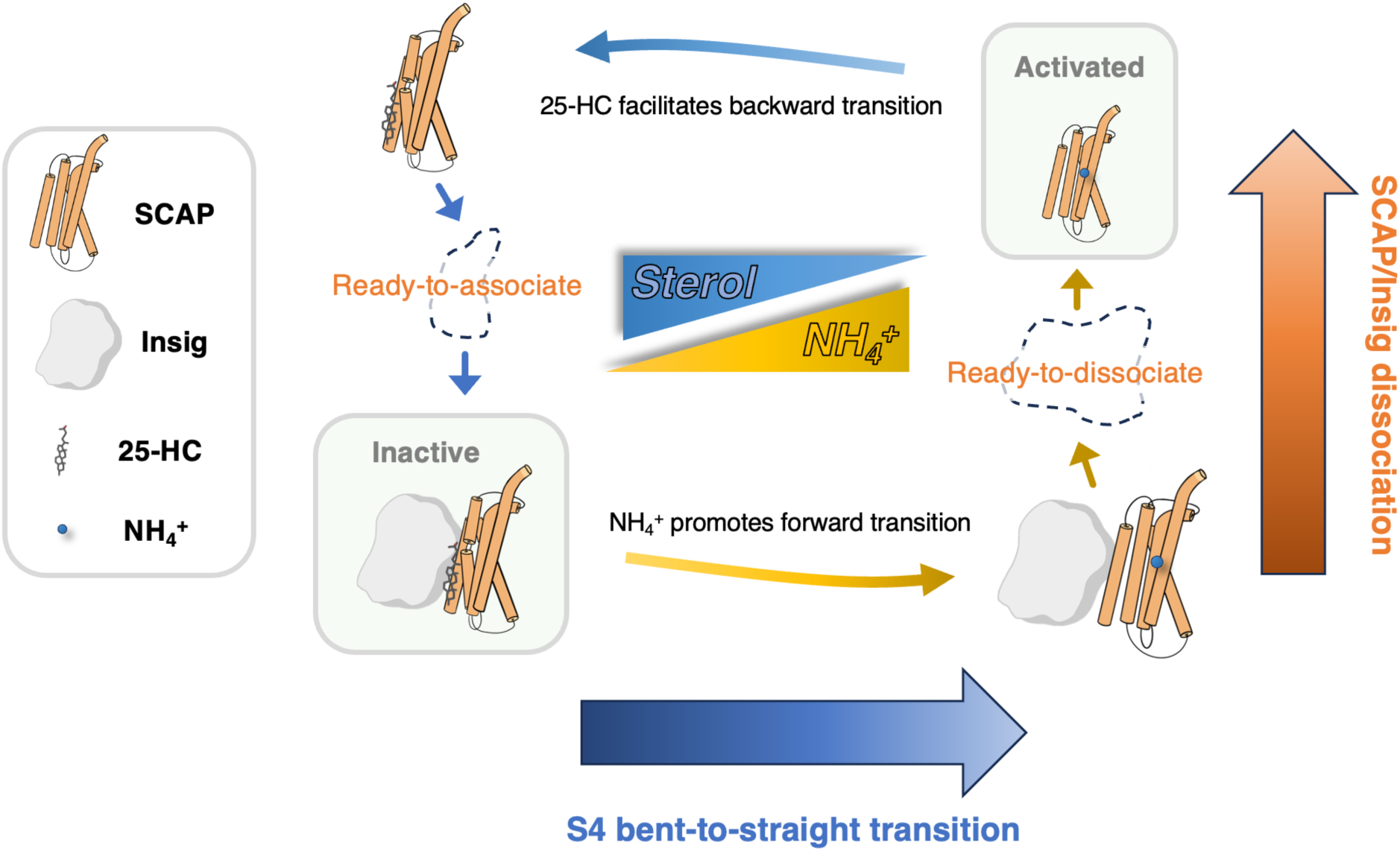
Schematic representation of the two key molecular events in SCAP activation: SCAP S4 conformational transition and SCAP-Insig association/dissociation, regulated by 25-HC and NH_4_^+^.

We next analyzed the hydrogen bonds at the SCAP-Insig interface (Figure 6c). In the inactive MD3, we identified two stable hydrogen bonds: one between Lys124 on the TM3-TM4 loop of Insig and Glu347 on the S4 helix of SCAP, and the other between Glu104 on the TM3 of Insig and Lys305 on the S2 helix of SCAP (Figure 6d). Additionally, the side chain of Arg110 on the TM3 of Insig oscillates between the side chains of Glu359 on the S4 helix and Glu399 on the S5 helix of SCAP, forming two dynamic hydrogen bonds. While the stable hydrogen bonds persisted in systems with a bent S4 conformation, the dynamic ones were less consistent (Figure S7). In contrast, the active systems featured only one stable hydrogen bond between Glu104 and Lys305 (Figure 6e). A decrease in the number of hydrogen bonds may indicate weaker interactions between the two proteins. These results highlight a direct correlation between the S4 conformational transition and the SCAP/Insig interaction, indicating the role of the unfolded S4 in coordinating the residue interaction network between SCAP and Insig.^4^ Our simulations further suggested that NH_4_^+^ or 25-HC does not directly stabilize the SCAP/Insig complex; instead, they modulate the SCAP/Insig interaction by selectively stabilizing either the bent or straight conformation of the S4 helix. Specifically, the depletion of 25-HC couples with NH_4_^+^ binding, facilitating the transition of the S4 helix from a bent to a straight conformation. This shift eventually promotes the dissociation of SCAP from Insig, as reflected by reduced interfacial contact area and polar contacts in the active system compared to the inactive one.

In our analysis of the structure and dynamics of the transmembrane helices of SCAP, we found a coupled motion between the S3 and S6 helices, with NH_4_^+^ serving as a hinge connecting these two helices (Figure S8a). Specifically, a pronounced tilting of the S6 helix, coupled with a subtler tilt in the S3 helix was observed only in the active system but not in the inactive one (Figure S8b). Additionally, these correlated helical rearrangements were absent when NH_4_^+^ bound to Gly357 on the S4 helix (MD1), rather than the bridging site formed by Asp428, Ser326 and Ser330 between the S3 and S6 helices (Figure S8c). Hence, both the NH_4_^+^ binding and the S4 conformational transition are essential to initiate the coupled motion between the S3 and S6 helices.

### Structure of the MELADL motif is allosterically regulated by NH4^+^ binding

A key step in SCAP activation is its translocation from the ER to the Golgi. This trafficking step relies on the recognition of SCAP by the COPII complex, specifically targeting the MELADL motif (residues 447 to 452). The MELADL motif is a hexapeptide sorting signal located on the cytoplasmic loop connecting the S6 and S7 helices of SCAP. Previous studies have unveiled that the association of SCAP with Insig can alter the spatial positioning of the MELADL motif relative to the membrane surface.^5^ Additionally, insertion or deletion of amino acid residues preceding the MELADL motif (between residues Thr437 and Arg446), presumably affecting its proximity to the ER membrane, were found to impede the trafficking of the SCAP/SREBP complex to the Golgi^5,37^. Interestingly, modifications following the MELADL motif did not seem to impact the translocation process.^5^ These findings suggest that the position of the MELADL motif relative to the membrane surface plays a crucial role in the recognition of SCAP by COPII. In light of this, we hypothesize that the depletion of 25-HC, coupled with NH_4_^+^ binding, not only stimulates the dissociation of SCAP from Insig, but may also enhance the accessibility of the MELADL motif to COPII.

We investigated the influence of alterations in the S6 helix orientation on the structure and spatial positioning of the MELADL motif relative to the membrane. In simulations of the active state of SCAP (MD2), the MELADL motif predominantly adopted an unstructured coil conformation, positioning it close to the membrane surface. This was evident from a decrease in the helix-to-membrane distance (Figure 7a). Conversely, in simulations of the inactive state (MD3), the MELADL motif maintained a helical conformation in the solvent throughout the microsecond-long simulations. This observation is corroborated by our radius of gyration analysis, which indicates a higher propensity for an unstructured MELADL motif in the active state (Figure S9).

While the precise molecular mechanism underlying the recognition of the MELADL motif remains largely unknown, we employed AF2 multimer to predict how the motif may interact with the membrane-facing side of Sec24. For human Sec24 subtypes A and B, both structured (helical) and unstructured forms of the MELADL were predicted to bind in the signaling peptide-binding pocket, as previously identified in yeast Sec24 (Figure 7b). This prediction aligns with our phylogenetic analyses (Figure S10a), illustrating a close relationship between Sec24a and Sec24b and their yeast counterparts. However, for more distantly related subtypes such as Sec24c, Sec24d, and Sec23, the model suggests the motif binding on the cytoplasmic side (Figure S10b).

Nevertheless, superimposition of our simulated MELADL conformations with the AF2-predicted SCAP-Sec24 complex structure suggests that a helical MELADL motif, as observed in the inactive SCAP, could impede its recognition by Sec24. This is due to potential clashes with the COPII protein assembly, including Sec24, Sec23, and Sar1 (Figure S11). Our observation of an unstructured yet stabilized MELADL motif on the membrane surface implies the potential importance of local membrane environments in facilitating this recognition.

## Conclusion

SREBPs are key transcription factors that regulate lipid metabolism through the SREBP pathway. Typically bound to SCAP, SREBPs are retained in the ER membrane. SREBP activation hinges on the dissociation of the SCAP/Insig complex, triggering the subsequent trafficking of the SCAP/SREBP complex to the Golgi and activating genes regulating lipid synthesis and uptake. Beyond factors like low cholesterol levels and SCAP N-glycosylation, our recent studies have identified ammonia as a key player in promoting the dissociation of N-glycosylated SCAP from Insig. In this study, extensive MD simulations were employed to unravel the structural mechanism through which ammonium ion binding activates SCAP. The SCAP S4 helix is believed to undergo a dynamic interconversion between two conformations: a partially unfolded, bent state and a fully intact, straight state. Our findings reveal a competitive binding between ammonium and 25-HC, with ammonium exhibiting a stronger affinity for the S4 helix in the straight conformation compared to the bent form. TMD simulations further suggest that ammonium acts as an agonist of SCAP, while 25-HC functions as an antagonist, stabilizing the active and inactive states of SCAP, respectively (Figure 8). Remarkably, ammonium operates as a hinge connecting the S3 and S6 helices, facilitating their coordinated movement. This hinge is critical for propagating the conformational transition of the S4 helix throughout the SCAP sterol sensing domain, inducing alterations in the orientation and structure of the S3 and S6 helices. These structural changes result in a decreased interaction interface contact area and a repositioned MELADL motif at the membrane surface.

In conclusion, our simulations unravel a detailed structural mechanism for ammonium-mediated SCAP activation. This mechanism delineates a sequence of early events integral to SCAP-Insig dissociation and MELADL recognition. Specifically, sterol depletion triggers conformational changes in the S4 helix, which, coupled with NH_4_^+^ binding, induces tilting motions between the transmembrane helices S3 and S6. These structural rearrangements not only facilitate the dissociation of SCAP from Insig, but also alter the exposure of the C-terminus of S6. The latter alteration influences both the position and structure of its sorting signal - the MELADL motif, potentially enhancing its accessibility to COPII, and thereby facilitating the recognition of SCAP by COPII to trigger the translocation of SCAP/SREBP from the ER to the Golgi. Our findings reveal the intricate interplay between 25-HC and ammonium binding and the conformational transition of the SCAP S4 helix, underscoring the pivotal role of the S4 helix in SCAP activation.

## Acknowledgements

This research was partially supported by The Ohio State University’s Translational Data Analytics Institute (TDAI) Interdisciplinary Research Pilot Award and used the resources of Ohio Supercomputer Center (OSC). K.C.C. also acknowledges the support from XJTLU startup fund and HPC platform.

