## Supplementary information for "Ammonia-mediated Conformational Dynamics in SCAP for SREBP Activation"

| CSM System | Co-solvent | 25HC-bound | S4 Conformation | Insig-bound | # of replicas |
|---|---|---|---|---|---|
| 1 | $NH_4^+$ | Yes | Bent | Yes | 5 |
| 2 | $NH_4^+$ | No | Bent | Yes | 5 |
| 3 | $NH_4^+$ | No | Straight | Yes | 5 |
| 4 | $NH_4^+$ | No | Straight | No | 5 |
| 5 | $NH_3$ | Yes | Bent | Yes | 5 |
| 6 | $NH_3$ | No | Bent | Yes | 5 |

**Table S1.** Summary of the CSM Simulation Systems.

| MD System | 25HC-bound | $NH_4^+$-bound | S4 Conformation | Insig-bound | # of replicas | Simulation times (μs) |
|---|---|---|---|---|---|---|
| 1 | No | Yes | Bent | Yes | 10 | 29.7 |
| 2 | No | Yes | Straight | Yes | 10 | 30.2 |
| 3 | Yes | No | Bent | Yes | 3 | 3.0 |
| 4 | No | No | Straight | Yes | 3 | 3.0 |
| 5 | No | No | Bent | No | 3 | 3.0 |

**Table S2.** Summary of the five MD Simulation Systems.

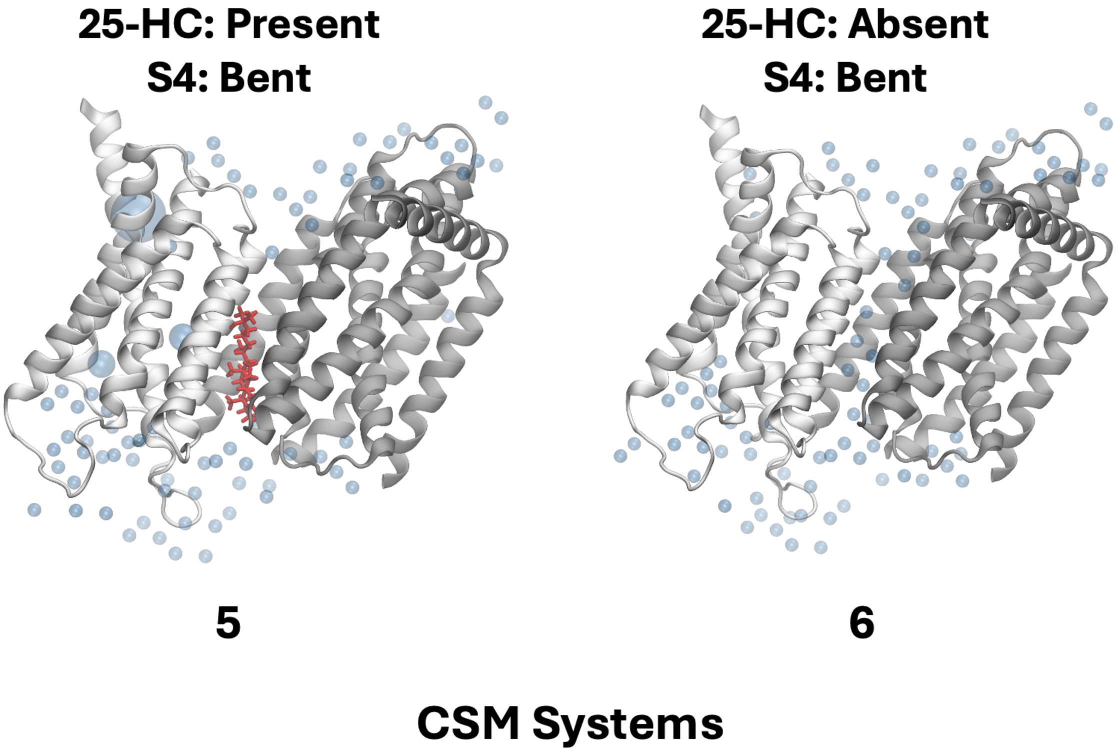


**Figure S1.** Distributions of co-solvent $NH_3$ near the SCAP-Insig complex. Blue spheres represent the locations of $NH_3$ molecules, with their radii proportional to the density averaged over the simulation trajectories.

Inactive Bent —Forward→ Active Straight

| Force constant | Apo | $NH_4^+$ bound | Y298C | D428A |
|---|---|---|---|---|
| 35 | | | 0.10 (30) | |
| 40 | | | 0.23 (30) | |
| 45 | | | 0.10 (30) | |
| 50 | 0.13 (10) | 0.30 (20) | 0.28 (40) | 0.22 (60) |
| 55 | 0.10 (20) | 0.80 (20) | 0.13 (30) | 0.33 (60) |
| 60 | | | | 0.50 (50) |
| 65 | 0.50 (30) | 0.60 (20) | 0.23 (20) | 0.63 (40) |
| 75 | 0.50 (40) | 0.90 (20) | 0.90 (20) | 0.70 (30) |
| 80 | | | 0.25 (20) | 0.94 (30) |
| 85 | 0.77 (43) | | | 0.70 (20) |
| 100 | 0.78 (40) | 1.00 (20) | 0.43 (23) | 0.83 (20) |
| 115 | 0.90 (20) | | | 0.90 (10) |
| 125 | 1.00 (10) | 1.00 (20) | 0.30 (20) | 1.00 (10) |
| 130 | 1.00 (10) | | | 0.90 (10) |
| 150 | 1.00 (10) | 1.00 (20) | | 0.80 (10) |

Probability

Inactive Bent ←Backward— Active Straight

| Force constant | Apo | $NH_4^+$ bound | Y298C | D428A |
|---|---|---|---|---|
| 20 | 0.10 (20) | | | 0.10 (20) |
| 25 | | | 0.90 (20) | |
| 30 | 0.60 (20) | 0.10 (20) | | 0.40 (20) |
| 35 | | | 0.70 (20) | |
| 40 | 1.00 (20) | 0.80 (20) | | 0.90 (20) |
| 45 | | | 0.50 (20) | |
| 50 | 1.00 (20) | 1.00 (20) | 0.85 (20) | 1.00 (20) |
| 75 | | | 0.38 (20) | |
| 80 | 1.00 (20) | 1.00 (20) | | 1.00 (20) |
| 100 | | | 0.36 (60) | |
| 120 | 1.00 (20) | 1.00 (20) | | 1.00 (20) |
| 125 | | | 0.30 (60) | |
| 150 | 1.00 (20) | 1.00 (20) | 0.34 (60) | 1.00 (20) |

Probability

**Figure S2.** Summary of probabilities for both forward and backward transitions under various simulation conditions. Number of replicas are shown in the parentheses.

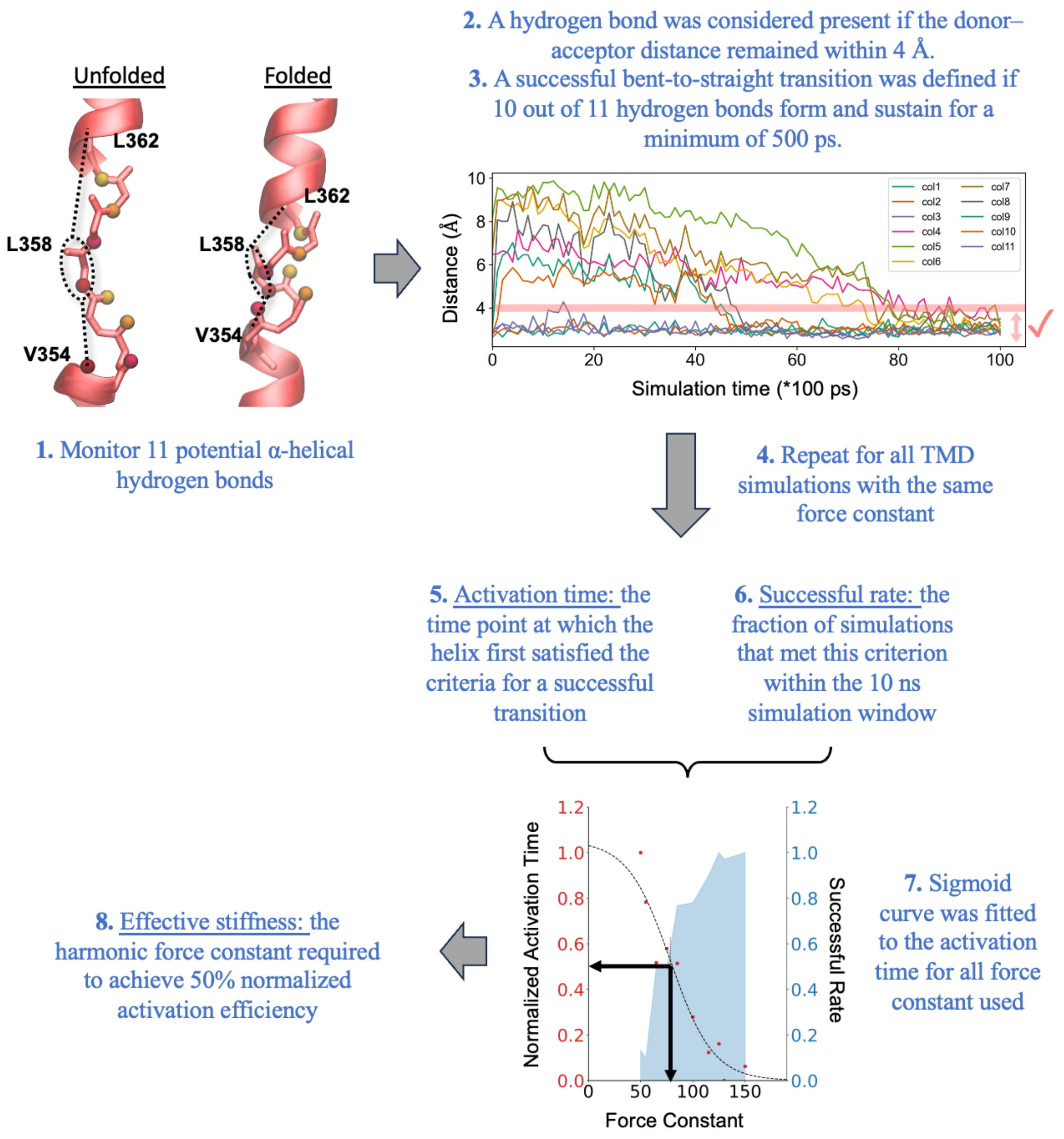


**Figure S3.** Schematic illustration of the methodology used to compute the effective stiffness of the SCAP structure, based on the ease of conformational transition between the bent (unfolded) and straight (folded) states of the S4 helix.

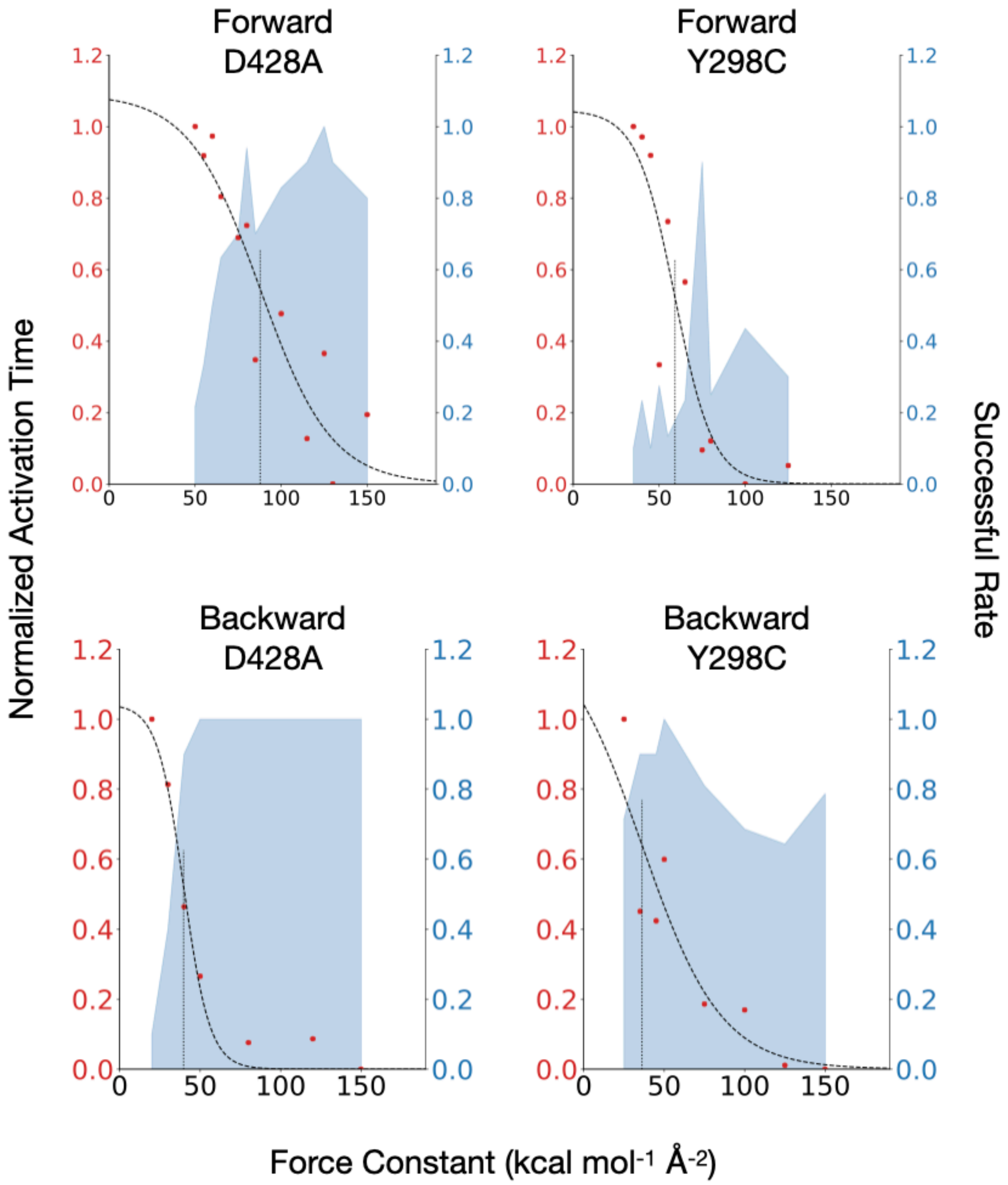


**Figure S4.** Activation times and success rates as a function of the TMD force constant for the forward and backward transition of the S4 conformation in two mutant systems (D428A and Y298C).

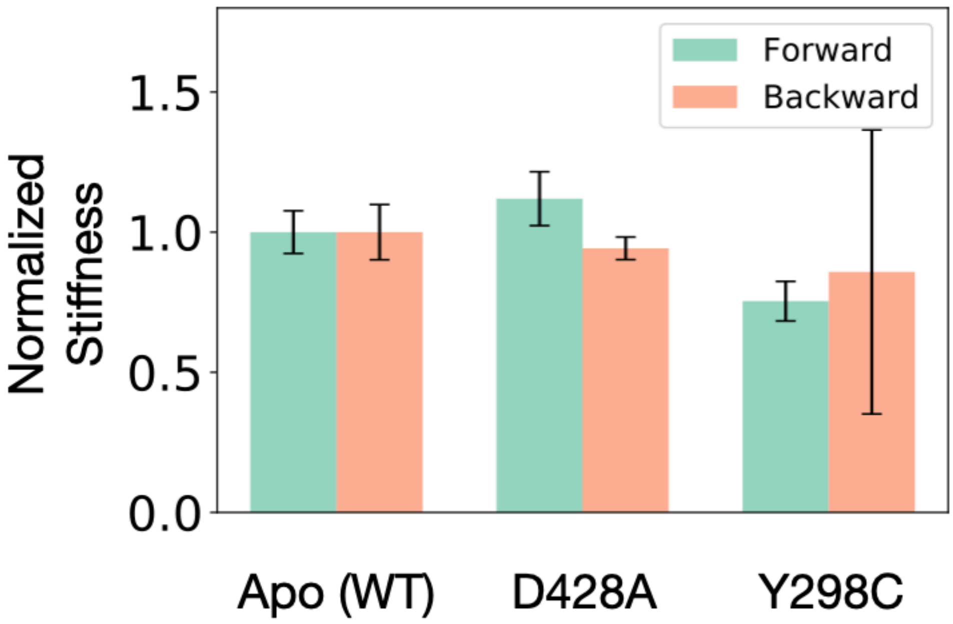


**Figure S5.** Normalized stiffness during the forward and backward transitions of the S4 conformation for the wild type and two mutant SCAP.

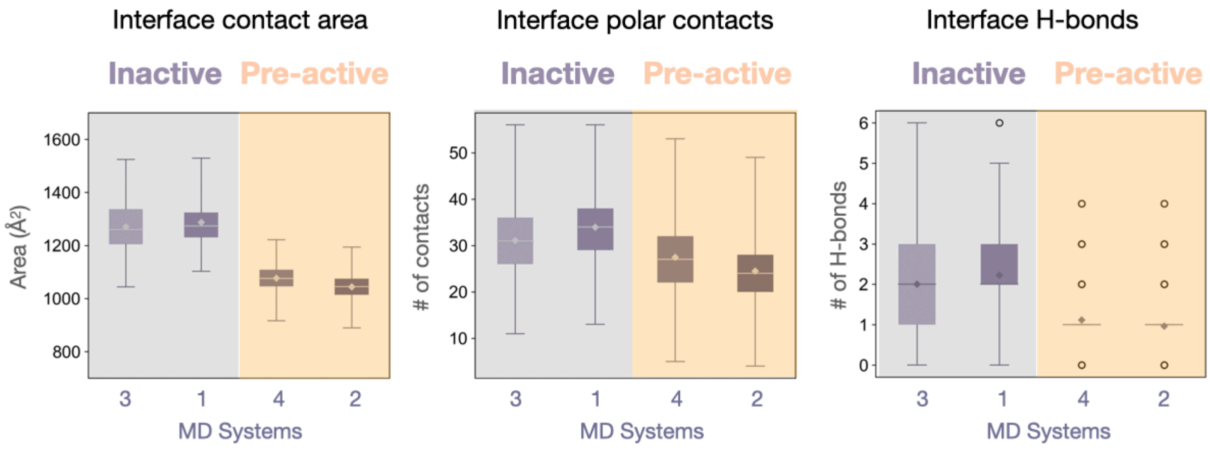


**Figure S6.** Comparison of interfacial contact areas, polar contacts, and hydrogen bonds across the four MD simulations of the inactive and pre-active systems.

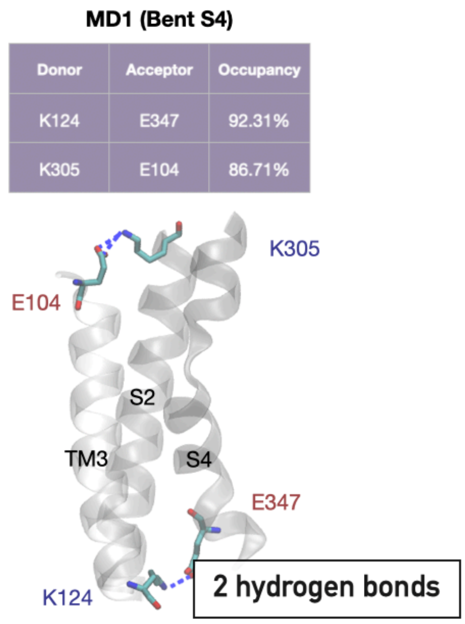


**Figure S7.** Interfacial hydrogen bonds between SCAP and Insig during the simulations of two presumably intermediate systems (MD1 and MD4).

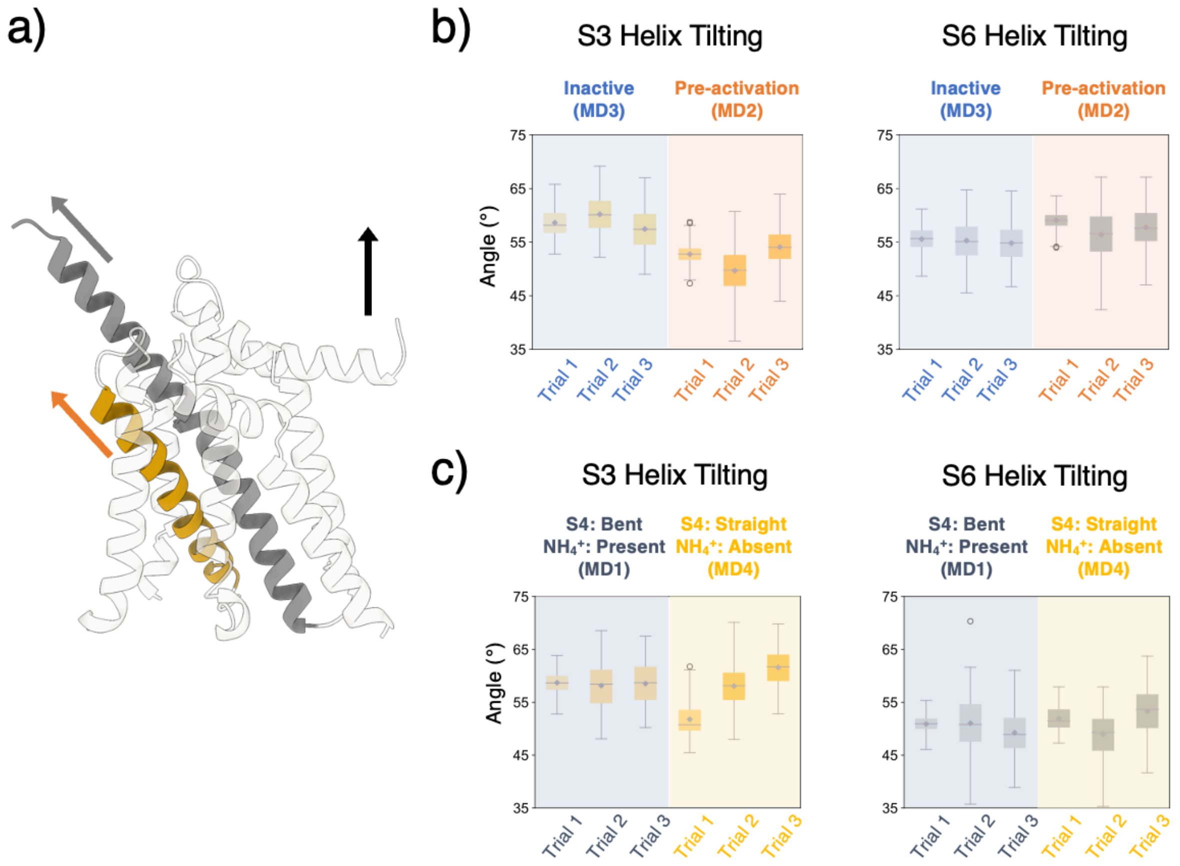


**Figure S8.** Comparative structural analysis of helical movement in inactive and active SCAP. (a) schematic representation of the relative tilting motion of helix S3 (colored in orange) and S6 (colored in grey) relative to the membrane normal (black arrow). (b) Comparison of the tilting angles of S3 and S6 helices in simulations of inactive (MD3) and pre-active (MD1) SCAP. (c) Comparison of the tilting angles of S3 and S6 helices in MD1 and MD4 simulations.

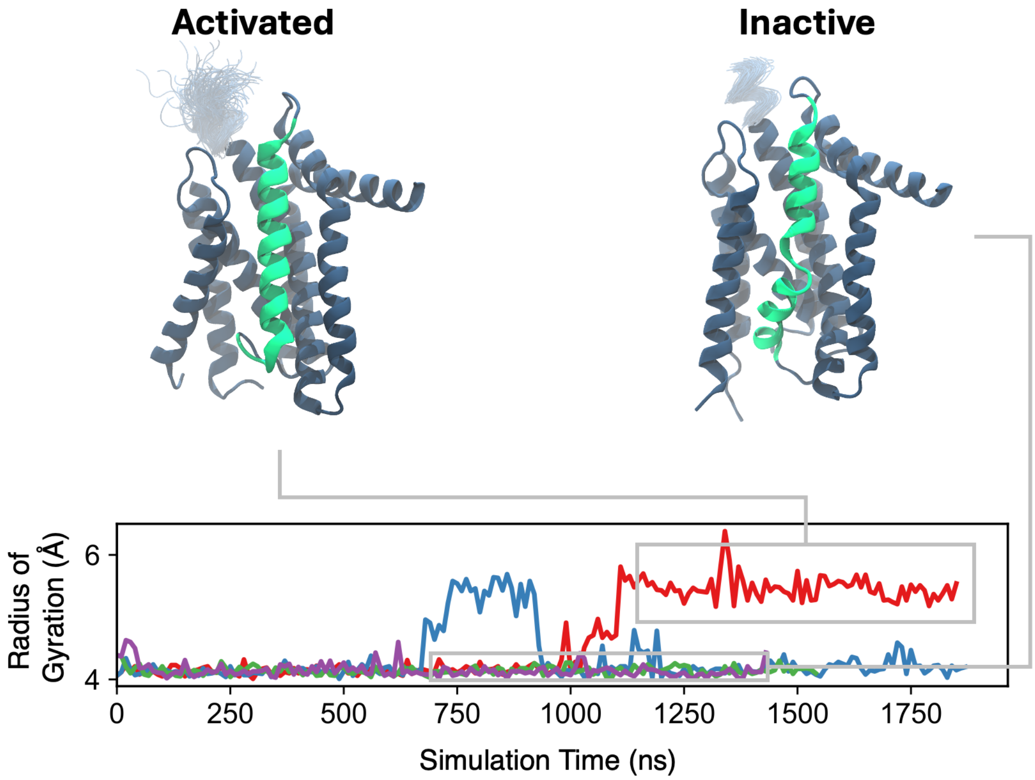


**Figure S9.** Radius of gyration of the C-terminus of the S6 helix as a function of time during the simulations of active (MD2) and inactive (MD3) SCAP.

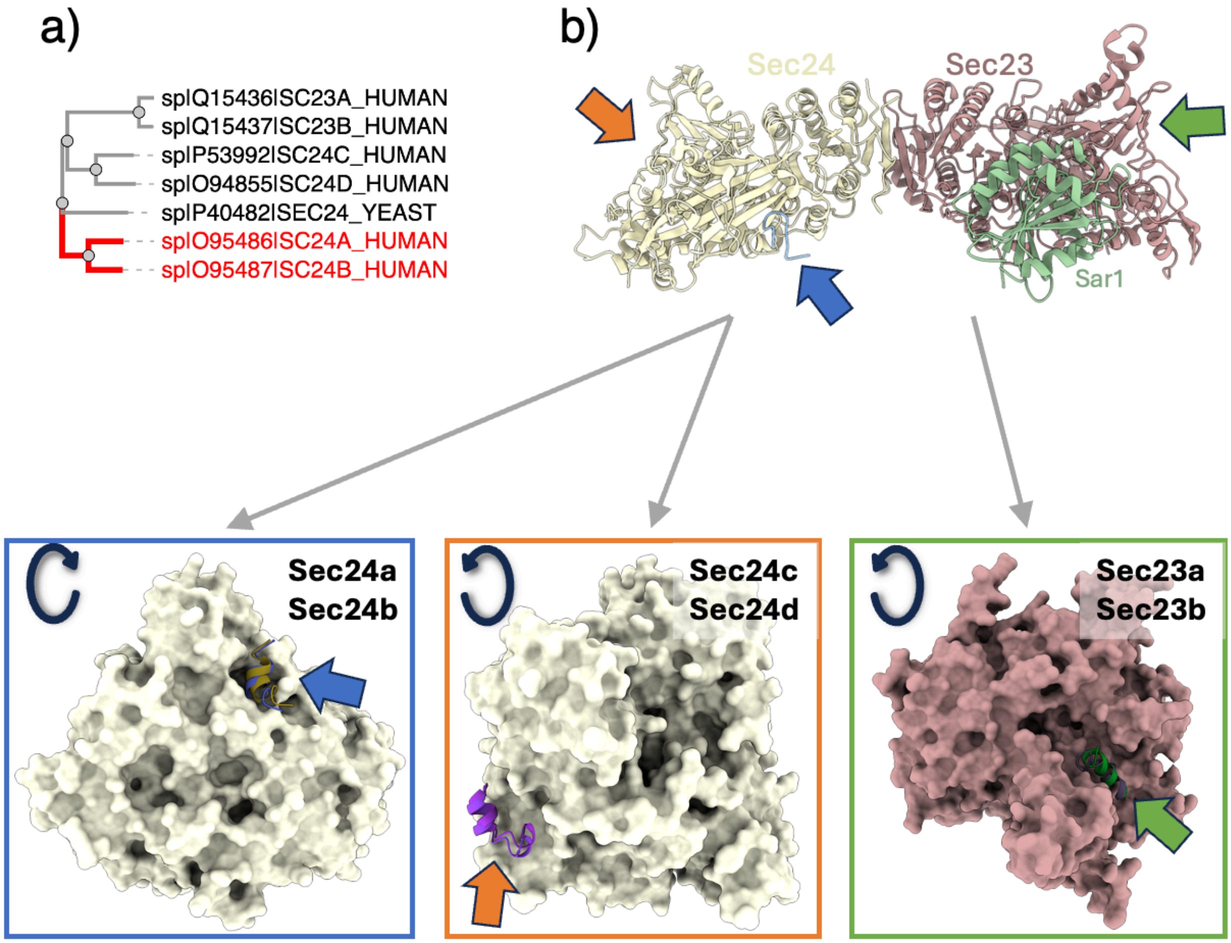


**Figure S10.** Binding of the MELADL motif to Sec proteins. (a) Phylogenetic analysis showcasing the evolutionary relationships among different Sec proteins. (b) Depiction of the binding sites of the MELADL motif on different Sec proteins.

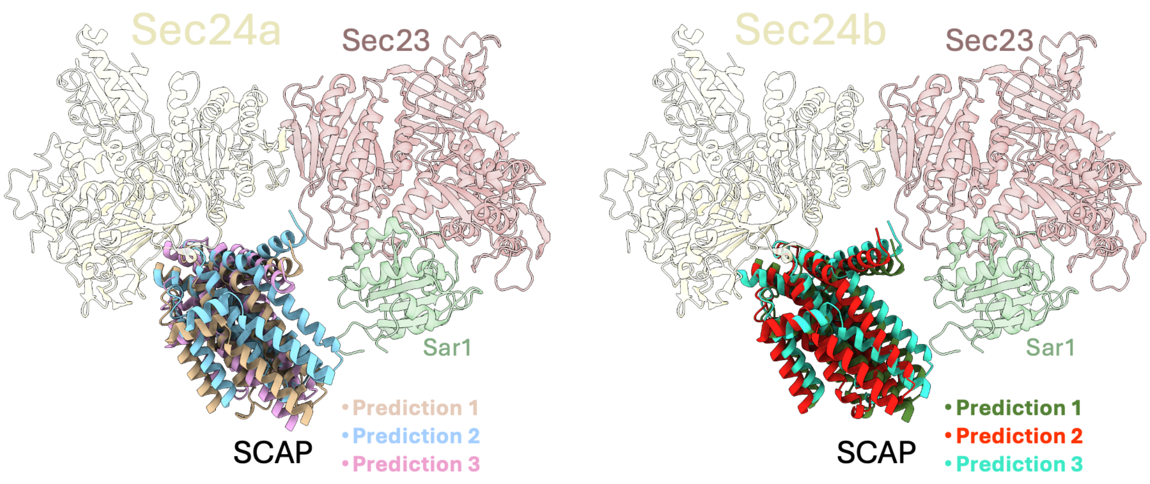


**Figure S11.** Structures of SCAP binding to the (left) Sec24a and (right) Sec24b complex predicted by AF2.
